# Cortical tracking of lexical speech units in a multi-talker background is immature in school-aged children

**DOI:** 10.1101/2022.04.29.490006

**Authors:** Maxime Niesen, Mathieu Bourguignon, Julie Bertels, Marc Vander Ghinst, Vincent Wens, Serge Goldman, Xavier De Tiege

## Abstract

Children have more difficulty perceiving speech in noise than adults. Whether these difficulties relate to immature processing of prosodic or linguistic elements of the attended speech is still unclear. To address the impact of noise on linguistic processing *per se*, we assessed how acoustic noise impacts the cortical tracking of intelligible speech devoid of prosody in school-aged children and adults.

Twenty adults and twenty children (7-9 years) listened to synthesized French monosyllabic words presented at 2.5 Hz, either randomly or in 4-word hierarchical structures wherein 2 words formed a phrase, and 2 phrases formed a sentence, with or without babble noise. Neuromagnetic responses to words, phrases and sentences were identified and source-localized.

Children and adults displayed significant cortical tracking of words in all conditions, and of phrases and sentences only when words formed meaningful sentences. In children compared with adults, cortical tracking of linguistic units was lower for all units in conditions without noise, and similarly impacted by the addition of babble noise for phrase and sentence units. Critically, when there was noise, adults increased the cortical tracking of monosyllabic words in the inferior frontal gyri but children did not.

This study demonstrates that the difficulties of school-aged children in understanding speech in a multi-talker background might be partly due to an immature identification of lexical but not supra-lexical linguistic units.

**Highlights:**

- Children track the hierarchical linguistic units of clear speech devoid of prosody
- This cortical tracking is left-hemisphere dominant as the adult brain
- Babble noise reduces cortical tracking of sentences in children and adults
- Unlike adults, children are not able to enhance cortical tracking of words in noise

## 1. Introduction

In daily life, humans tend to gather in places often unfavorable to verbal communication due to noisy backgrounds. In such situations, successful conversation is challenging since listeners have to tune in to the speaker’s voice, while tuning out the noisy auditory scene. This difficult task is especially challenging for children, who typically have lower speech in noise (SiN) processing abilities than adults (Elliott, 1979; Hall et al., 2002). Whether these lower abilities relate to a higher impact of noise on the neural processing of speech linguistic or paralinguistic (e.g., prosody) information in children is still unsettled.

One way to understand how the human brain processes SiN is to study how cortical activity tracks the fluctuations of natural connected speech in noisy backgrounds. Such cortical tracking of speech (CTS) typically occurs at frequencies matching with the hierarchical temporal linguistic units (i.e., syllables, words, phrases/sentences) and with paralinguistic information such as prosodic cues (Giraud and Poeppel, 2012; Gross et al., 2013; Ding and Simon, 2014). It is considered to subserve the incremental neural grouping of abstract linguistic units to promote subsequent speech recognition (Ding et al., 2016). In quiet background, CTS is observed in school-aged children but is reduced at the syllabic level compared with adults (Vander Ghinst et al., 2019; Bertels et al., 2022). In multi-talker situations, their auditory system selectively tracks the attended speech stream rather than the global auditory scene. Still, CTS in children is more compromised by increasing noise level for words and phrases/sentences compared with CTS in adults. Furthermore, syllabic CTS in children does not increase in babble noise as observed in adults (Vander Ghinst et al., 2019; Bertels et al., 2022).

Still unclear is whether the higher impact of noise on children’s CTS relates to increased alterations in the cortical tracking of the attended hierarchical linguistic units, or of the paralinguistic information such as prosody. Indeed, previous studies used natural connected speech masked by babble noise, yet natural connected speech comprises both hierarchical linguistic and paralinguistic information (e.g., prosody) that temporally correlate (Yamashita, 2013). One way to specifically address the impact of noise on the neural processing of speech linguistic units is to eliminate prosodic cues from the attended speech stream. In quiet environments, such an approach has already evidenced successful cortical tracking of hierarchical linguistic units in adults (Yamashita, 2013; Ding et al., 2016, 2017). In this context, the adults’ brain internally groups small linguistic units (words) into larger linguistic units (phrases and sentences), based on grammar knowledge only (Ding et al., 2016, 2017).

In the present study, we used speech devoid of prosody to contrast grammar-based CTS between children and adults, and between quiet and noisy conditions. To this end, we quantified using magnetoencephalography (MEG) the cortical tracking of isochronous monosyllabic words either forming or not phrases/sentences in the absence or presence of babble noise. As grammar-based tracking requires increased attention (Makov et al., 2017; Ding et al., 2018) and becomes less accurate when speech intelligibility decreases (Blanco-Elorrieta et al., 2020; Meng et al., 2021), we hypothesized that babble noise would impede the grammar-based internal grouping of monosyllabic words into phrases/sentences. Furthermore, as syntactic and semantic processing are developing until late childhood (Nuñez et al., 2011; Skeide and Friederici, 2016) and as children have lower SiN abilities than adults, we also hypothesized that grammar-based tracking of speech would be affected by babble noise to a greater extent in children compared with adults. Finally, we also hypothesized that the cortical tracking of small linguistic units such as monosyllabic words would be differently impacted by babble noise in children and adults, as previously shown with natural connected speech (Vander Ghinst et al., 2019; Bertels et al., 2022).

## 2. Methods

### 2.1 Subjects

Twenty healthy adults (mean age: 24 years, age range: 20–29 years, 11 females) and twenty healthy children (mean age: 8 years, age range: 7–9 years, 9 females) took part in this study. All subjects were native French speakers and right-handed according to the Edinburgh Handedness Inventory (Oldfield, 1971) (adults score range: 50–95, mean score: 75; children score range: 50–95, mean score: 73.5; *t_38_* = 0.30, *p* = 0.77). They all had no history of neurological, psychiatric or otologic disorders and had normal hearing according to pure tone audiometry (i.e., normal hearing thresholds: between 0–20 dB HL for 125 Hz to 8000 Hz). Using three separate subtests (a speech audiometry, a SiN audiometry and a dichotic test) of a validated and standardized central auditory battery in French (Demanez et al., 2003), we have ensured that participants had normal dichotic and SiN perception for their age.

This study was approved by the Ethics Committee of the HUB-Hôpital Erasme. All subjects (and their legal representatives for children) gave written informed consent prior to their inclusion in the study.

### 2.2 Stimuli

Figure 1 illustrates the stimuli used in the present study. The stimuli were adapted from those described in (Ding et al., 2016) to the French language.

**Figure 1.**
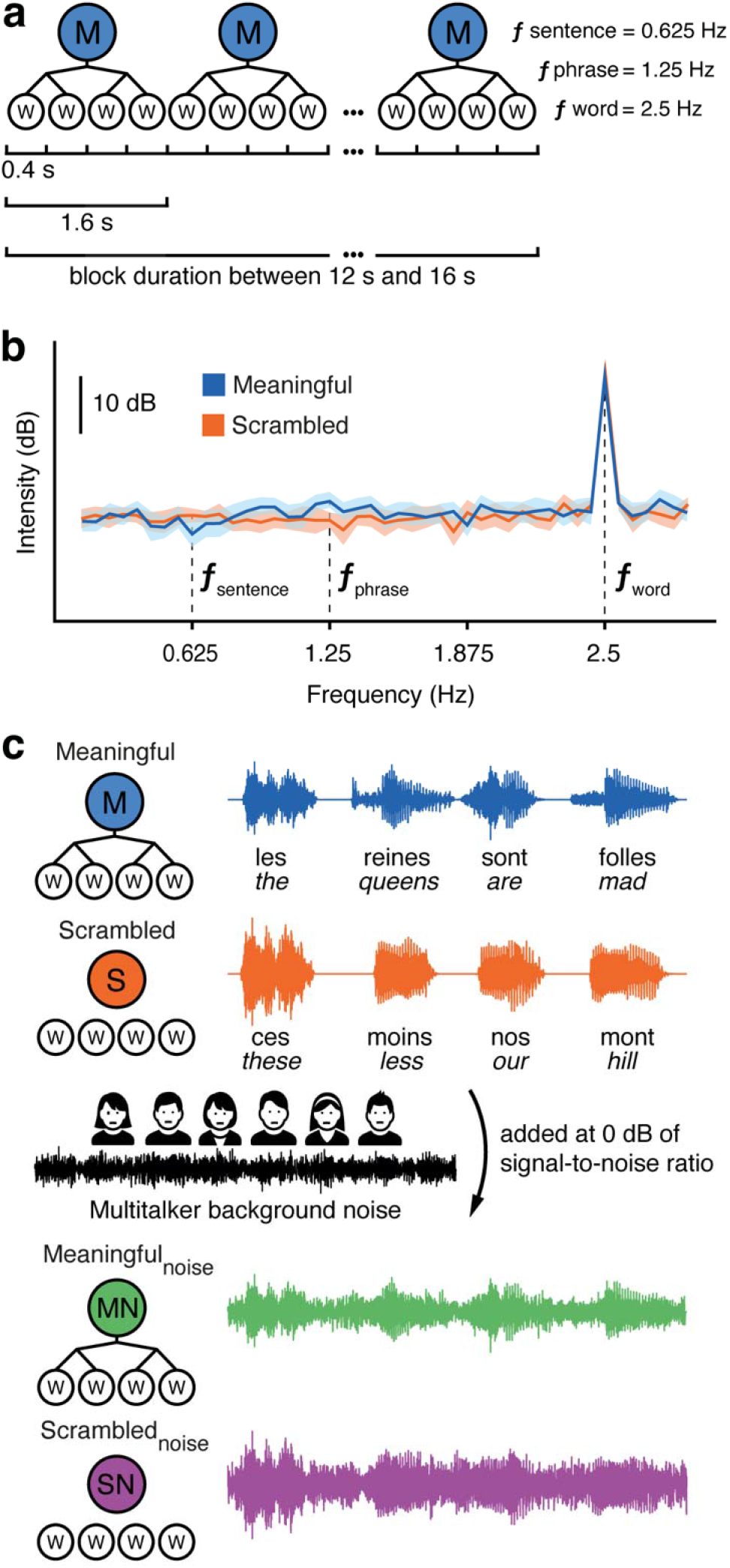
Experimental stimuli. **(a)** Sequence of monosyllabic words (presentation rate 2.5 Hz) forming phrases (1.25 Hz, determiner + noun and verb + adjective/adverb) and sentences (0.625 Hz, determiner + noun + verb + adjective/adverb). **(b)** Spectrum of stimulus intensity disclosing a clear peak at the monosyllabic word rate but not at the phrase or sentence rates. **(c)** Time-course of stimuli for each condition.

The auditory stimuli were 238 different monosyllabic French words. They were synthesized using the MacinTalk Synthesizer (male voice, Thomas, in macOs Sierra 10.12.6) and were adjusted to the same intensity and the same duration of 400 ms by truncation (without alteration of word identity) or silence padding symmetrically on both sides (original mean duration of 375 ± 70 ms, range 146–525 ms). To introduce a fade-in and a fade-out, the extremities of each word were multiplied by a 25-ms squared-sine ramp signal.

The stimuli were used to build blocks of 40 words in 4 different conditions described hereafter (*Meaningful, Meaningful_noise_, Scrambled*, and *Scrambled_noise_*). Word acoustic waveforms were concatenated without any additional acoustic gap between words. Considering the duration of each word (i.e., 400 ms), the word rate was 2.5 Hz (*f*_word_). Twenty-five different blocks were built for each condition.

For the *Meaningful* condition, we constructed 250 different sentences composed of four monosyllabic words. All sentences shared the same hierarchical linguistic structures: determiner + noun + verb + adjective/adverb. In that setting, the phrase (i.e., determiner + noun and verb + adjective/adverb) rate was 1.25 Hz (*f*_phrase_), and the sentence (determiner + noun + verb + adjective/adverb) rate was 0.625 Hz (*f*_sentence_) (**Figure 1a)**. Critically, as in (Ding et al., 2016), the linguistic units could only be extracted using grammar-based knowledge, and not prosodic cues. Indeed, the sound envelope featured fluctuations at *f*_word_ but not at *f*_phrase_ or *f*_sentence_ due to the absence of prosodic cues for phrase/sentence boundaries (**Figure 1b)**.

The *Scrambled* condition was created by randomly shuffling the order of the monosyllabic words used in the *Meaningful* condition, resulting in meaningless strings of words presented at 2.5 Hz.

Two more listening conditions, *Meaningful_noise_* and *Scrambled_noise_*, were created by adding a multitalker background noise to the *Meaningful* and *Scrambled* conditions at a signal-to-noise ratio (SNR) of 0 dB (**Figure 1c**). This SNR was chosen because it is typically encountered in multi-talker situations (Bronkhorst, 2015). The background noise was already used and described in previous studies from our group (Vander Ghinst et al., 2016, 2019). Briefly, it consisted of a mix of 6 native French speakers’ speech (3 females and 3 males). This configuration was chosen because it introduces interference at phonetic and lexical levels (Simpson and Cooke, 2005; Hoen et al., 2007).

### 2.3 Experimental paradigm

During MEG recordings, subjects sat comfortably in the MEG chair with their arms laying on a table. They were asked to gaze at a cross on the wall in front of them and not to move. After a 5-min rest condition (i.e., stimulation-free), subjects underwent five listening sessions, each lasting ~6 min. Listening sessions consisted of 20 blocks randomly selected among the four different conditions (i.e., *Meaningful, Scrambled, Meaningful_noise_, Scrambled_noise_*) with the rule that two consecutive blocks cannot be of the same condition. The order of blocks was randomized across conditions and blocks were separated by a silent break of 3 s.

Auditory stimulations were played using VLC Media Player (VideoLAN Project, GNU General Public License) and were delivered through a MEG-compatible 60 × 60 cm^2^ high-quality flat panel loudspeaker (Panphonics SSH sound shower, Panphonics Oy) placed ~2.5 m in front of the subjects. The average sound intensity was set to 60 dB as assessed by a sound level meter (Sphynx Audio System).

To ensure subjects maintained their attentional focus on the auditory stimuli, they were asked to repeat the last word they heard at the end of each block during the acoustic gap (a behavioral task henceforth referred to as “word identification”). To avoid a possible prediction of the word to be repeated based on the preceding linguistic units, we randomly truncated each block by a number of words in between 0 and 8, leading to blocks of 32–40 words. The last word to be repeated could therefore be of any class. After the MEG recordings, subjects were asked to rate the intelligibility of a randomly chosen block of each condition on a visual analog scale ranging from 0 to 10 (0, totally unintelligible; 10, perfectly intelligible).

### 2.4 Data acquisition

Neuromagnetic signals were recorded at the HUB-Hôpital Erasme with a whole-scalp-covering MEG system (Triux, MEGIN, Croton Healthcare, Finland) installed in a lightweight magnetically shielded room (Maxshield, MEGIN, Croton Healthcare, Finland; see (De Tiège et al., 2008). The MEG system comprised 102 sensor triplets, each consisting of one magnetometer and two orthogonal planar gradiometers. The recording bandpass filter was 0.1–330 Hz and the data were sampled at 1 kHz. We used four head-tracking coils to continuously monitor subjects’ head position inside the MEG helmet. We digitized with an electromagnetic tracker (Fastrak, Polhemus) the location of the coils and at least 250 head-surface points (on scalp, nose, and face) with respect to anatomical fiducials.

Subjects’ high-resolution 3D-T1 weighted cerebral magnetic resonance images (MRI) were acquired on a 1.5 T MRI (Intera, Philips).

### 2.5 Data preprocessing

Continuous MEG data were first preprocessed off-line using the temporal signal space separation method implemented in MaxFilter software (MaxFilter, MEGIN, Croton Healthcare, Finland; correlation limit 0.9, segment length 20 s) to suppress external interferences and to correct for head movements (Taulu et al., 2005; Taulu and Simola, 2006).

To further suppress physiological artifacts from MEG data, 30 independent components were evaluated from the data band-pass filtered at 0.1–25 Hz and reduced to a rank of 30 with principal component analysis. Independent components corresponding to heartbeat, eye-blink, and eye-movement artifacts were visually identified, and corresponding MEG signals reconstructed by means of the mixing matrix were subtracted from the full-rank and full-band data. Across subjects and conditions, the number of components rejected was 2.5 ± 0.5 (mean ± SD) in the adult group and 3.8 ± 1.0 in the children group (*t_38_* = 5.06, *p_corr_* < 0.0001).

The resulting data were then filtered through 0.1–40 Hz using a zero phase-lag FFT filter. MEG epochs were extracted from the 5^th^ word onset (to avoid the transient response to the acoustic onset of each block) to the 32^nd^ word offset (because of the random truncation of blocks). Epochs were considered contaminated by artifacts and removed from further analyses when their maximum MEG amplitude exceeded 5 pT in at least one magnetometer or 1 pT/cm in at least one gradiometer. The mean ± SD number of artifact-free epochs was 24.3 ± 2.0 (across subjects and conditions) in the adult group and 22.2 ± 2.7 in the children group (*F_1,38_* = 4.64, *p* < 0.038). To avoid a possible methodological bias in our results due to differences between age groups in the number of epochs analyzed, we randomly discarded epochs in adults’ data to equalize the number of epochs in both groups.

### 2.6 Sensor-space data analyses

Retained epochs were Fourier-transformed (frequency resolution 0.089 Hz). For each subject, condition and sensor, amplitude spectra were obtained as the modulus of the averaged Fourier-transformed epochs. Note that because the modulus was taken after averaging Fourier coefficients, our derivation of amplitude spectra allowed for phase cancellation of activity not phase-locked with audio sequences. At each sensor triplet, we retained only the Euclidean norm of the amplitude across pairs of planar gradiometers. For each subject, condition and sensor, SNR responses were computed as the ratio between the amplitude at each frequency bin and the average amplitude at the 10 surrounding frequency bins (5 on each side, excluding the immediately adjacent bins) (Peykarjou et al., 2017; Barry-Anwar et al., 2018; Bertels et al., 2020). SNR values significantly above 1 at *f*_word_, *f*_phrase_ or *f*_sentence_ would indicate specific discrimination of the corresponding linguistic units.

### 2.7 Source-space data analyses

Source reconstruction was used to estimate brain maps of SNR. For that, MEG and MRI coordinate systems were co-registered using the 3 anatomical fiducial points for initial estimation and the head-surface points for further manual refinement. Then, the individual MRIs were segmented using Freesurfer software (Martinos Center for Biomedical Imaging, Boston, MA, RRID:SCR_001847; (Reuter et al., 2012)), and a non-linear transformation from individual MRIs to the MNI brain was computed using the spatial normalization algorithm implemented in Statistical Parametric Mapping (SPM8, Wellcome Department of Cognitive Neurology, London, UK, RRID:SCR_007037; (Ashburner et al., 1997; Ashburner and Friston, 1999)). This transformation was used to map a homogeneous 5-mm grid sampling the MNI brain volume onto individual brain volumes. For each subject and grid point, the MEG forward model corresponding to three orthogonal current dipoles was computed using the one-layer Boundary Element Method implemented in the MNE software suite (Martinos Center for Biomedical Imaging, Boston, MA, RRID:SCR_005972; (Gramfort et al., 2014)). The forward model was then reduced to its two first principal components. This procedure is justified by the insensitivity of MEG to currents radial to the skull, and hence, this dimension reduction leads to considering only the tangential sources. A Minimum-Norm Estimates inverse solution (Dale and Sereno, 1993) was then used to project sensor-level Fourier coefficients (averaged across epochs) into the source space. We followed the same approach as that used at the sensor level to estimate source-level SNR (source pairs taking the place of gradiometer pairs).

We further identified the coordinates of local maxima in group-averaged SNR maps. Such local maxima of SNR are sets of contiguous voxels displaying higher SNR values than all neighboring voxels. We only report statistically significant local maxima of SNR, disregarding the extent of these clusters. Indeed, cluster extent is hardly interpretable in view of the inherent smoothness of MEG source reconstruction (Hämäläinen and Ilmoniemi, 1994; Wens et al., 2015; Bourguignon et al., 2018).

The significant local maxima were visualized on the MNI glass brain using the BrainNet viewer (Xia et al., 2013) (see **Figure 5**).

### 2.8 Statistical analysis

#### 2.8.1 Behavioral results

Because results of speech and SiN audiometry were not normally distributed as indicated by Shapiro-Wilk tests (both *p* < 0.05 for children and adults’ results in silence), we performed a *Mann-Whitney U* test to compare and identify statistical differences between adults and children.

A three-way repeated-measures ANOVA was used to assess the effects of condition (within-subject factor; *Meaningful*, *Scrambled*, *Meaningful_noise_, Scrambled_noise_*), position (within-subject factor; determiner, noun, verb, adjective/adverb) and of age group (between-subjects factor; children, adults) on word identification.

A two-way repeated-measures ANOVA was used to assess the effects of condition (within-subject factor; *Meaningful*, *Scrambled*, *Meaningful_noise_, Scrambled_noise_*) and of age group (between-subjects factor, children, adults) on the intelligibility rating.

Post hoc comparisons were performed with pairwise t tests with Bonferroni adjustment for multiple comparisons.

#### 2.8.2 Individual levels of SNR

A nonparametric permutation-like test, first described in (Bertels et al., 2020), was used to estimate the statistical significance of the SNR (subject-level) at *f*_word_, *f*_phrase_ and *f*_sentence_ separately. The test sought for significant responses in all gradiometer pairs, with correction for multiple comparisons across them. Such a statistical test was chosen because it can support claims of statistical significance at each gradiometer pair separately, in contrast with, e.g., cluster-based permutation tests (Sassenhagen and Draschkow, 2019). In a nutshell, the statistical procedure trims the epochs to randomize the position of the linguistic units under assessment and hence to destroy the phase locking across epochs of possible responses specific to these units. The parameters of the test for phrase and sentence SNR were different from those for word SNR. We therefore present the procedure for phrase and sentence SNR, and highlight the modifications for word SNR.

To test the significance of the SNR at *f*_phrase_ and *f*_sentence_, a permutation distribution for that SNR was built by estimating 1000 times the maximum across gradiometer pairs of the SNR at the tested frequency bin derived from epochs randomly trimmed by a duration corresponding to the *n* = 0, 1, 2 or 3 first words (*n* × 400 ms) and 4–*n* last words. To match epoch length across permuted and genuine data, genuine SNR in each gradiometer pair was re-computed based on epochs in which either the first or last 1.6 s of data (corresponding to 4 words) was removed. The significance of the genuine response at each gradiometer pair was computed as the proportion of values in the permutation distribution that were above the observed genuine value. This test, being akin to a permutation test (Nichols and Holmes, 2002), is exact, and because the permutation distribution was built on maximum values across gradiometer pairs, it intrinsically deals with the multiple comparison issue.

The difference to test the significance at word frequencies laid in the trimming scheme. Word SNR was recomputed based on epochs in which either the first or last 400 ms of data (corresponding to 1 word) was removed; and to estimate the permutation distribution, epochs were randomly trimmed by a duration corresponding to the *n* = 0 or 1 first half-words (*n* × 200 ms) and 2–*n* last half-words. The trimming scheme randomized the position of words within epochs, so that epochs either started at word onset or in the middle of a word.

#### 2.8.3 Local maxima of group-level SNR

The statistical significance of the local maxima of SNR observed in group-averaged maps for each age group, condition and frequency of interest was assessed with a non-parametric permutation test that intrinsically corrects for multiple spatial comparisons (Nichols and Holmes, 2002). First, subject and group-averaged *rest* maps of SNR were computed with MEG epochs randomly extracted from the rest condition. Group-averaged difference maps were obtained by subtracting *genuine* and *rest* group-averaged SNR maps. Under the null hypothesis that SNR maps are the same whatever the experimental condition, the labeling *genuine* or *rest* are exchangeable prior to difference map computation (Nichols and Holmes, 2002). To reject this hypothesis and to compute a significance level for the correctly labeled difference map, the sample distribution of the maximum of the difference map’s absolute value within the entire brain was computed from a subset of 1000 permutations. The threshold at *p* < 0.05 was computed as the 95 percentile of the sample distribution (Nichols and Holmes, 2002). All supra-threshold local maxima of SNR were interpreted as indicative of brain regions showing statistically significant CTS and will be referred to as sources of CTS.

Permutation tests can be too conservative for voxels other than the one with the maximum observed statistic (Nichols and Holmes, 2002). For example, dominant SNR values in the right hemisphere could bias the permutation distribution and overshadow weaker SNR values in the left hemisphere, even if these were highly consistent across subjects. Therefore, the permutation test described above was conducted separately for left- and right-hemisphere voxels.

#### 2.8.4 Effect of age group and noise on SNR

We used repeated-measures ANOVAs to compare the SNR of sources of CTS between adults and children with an additional factor of noise (*Meaningful* and *Meaningful_noise_* conditions). The dependent variable was the maximum SNR value within a sphere of 10-mm radius around the maxima of the group-level SNR map averaged across age groups and conditions in order to limit potential bias coming from differences in source location. However, since all sources of CTS were bilateral, the factor hemisphere (left and right) was added to the ANOVA. Separate ANOVAs were run for word, phrase, and sentence SNR, and for the different local maxima. Post hoc comparisons were performed with pairwise t tests with Bonferroni adjustment for multiple comparisons.

### 3. Results

#### 3.1 Behavioral results

Speech audiometry in silence did not differ between adults and children (mean score ± SD: adults = 28.3 ± 0.9, children = 28.1 ± 1.1, *U* = 232.5, *p* = 0.356), but differed in noise (adults = 27.4 ± 1.2, children = 25.9 ± 2.0, *U* = 284.5, *p* = 0.020).

Similar results were observed for word identification (**Figure 2a**). Indeed, the ANOVA performed on these scores revealed a significant effect of age group (*F_1,38_* = 39.9, *p* < 0.0001), a significant effect of condition (*F_3,114_* = 200.8, *p* < 0.0001) and a significant interaction between age group and condition (*F_3,114_* = 8.7, *p* < 0.0001), but no effect of word class (determiner, noun, verb, adjective/adverb) in the sentence (*F_3,114_* < 1) nor interaction involving this latter factor (all *p* > 0.05). *Post-hoc* comparisons between conditions demonstrated that scores were (1) not significantly different between *Meaningful* and *Scrambled* conditions (adults, *t_19_* = 1.26, *p_corr_* = 1; children, *t_19_* = 2.23, *p_corr_* = 0.67) but significantly higher in noiseless conditions compared to noisy conditions (all *p_corr_* < 0.0001), (2) significantly higher in *Meaningful_noise_* compared with *Scrambled_noise_* condition (adults, *t_19_* = 8.64, *p_corr_* < 0.0001; children, *t_19_* = 5.57,*p_corr_* < 0.0001), and (3) lower in children compared with adults only in noisy conditions (*Meaningful_noise_, t_38_* = 7.11, *p_corr_* < 0.0001; *Scrambled_noise_, t_38_* = 4.11, *p_corr_* = 0.002).

**Figure 2.**
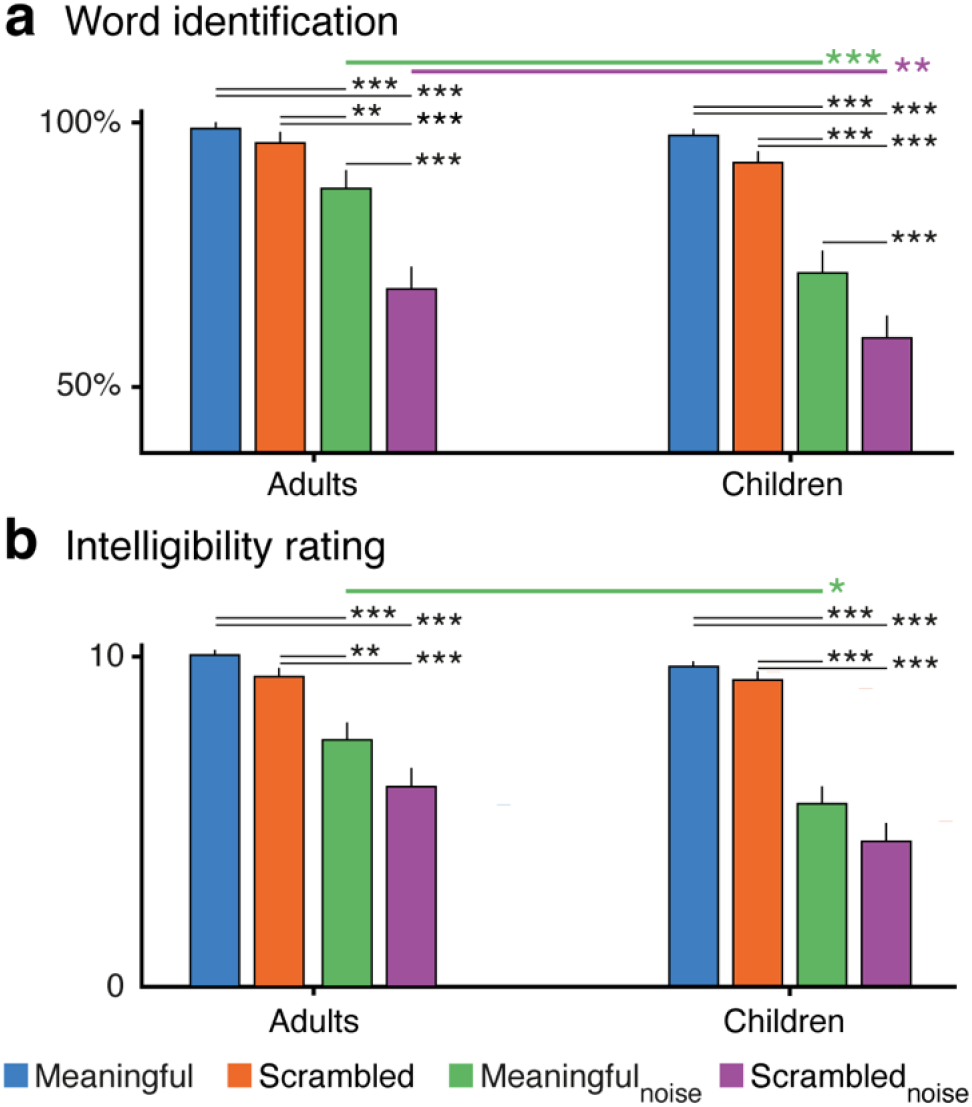
Mean ± SD scores for the word identification (**a**) and the intelligibility rating (**b**) in both age groups. Asterisks indicate significant differences (* = *p* < 0.05, ** = *p* < 0.01, *** = *p* < 0.001).

The ANOVA performed on the intelligibility ratings (**Figure 2b**) revealed the same significant effect of age group (*F_1,38_* = 5.99, *p_corr_* = 0.019) and condition (*F_3,114_* = 84.5, *p_corr_* < 0.0001), and a significant interaction between age group and condition (*F_3,114_* = 9.52, *p_corr_* = 0.009). *Post-hoc* comparisons demonstrated that intelligibility ratings were (1) significantly higher in noiseless compared to noisy conditions (all *p_corr_* < 0.01), (2) not significantly different between *Meaningful* and *Scrambled* conditions nor between *Meaningful_noise_* and *Scrambled_noise_* conditions (all *p_corr_* > 0.05), and (3) lower in children compared with adults only in the *Meaningful_noise_* condition (*t_38_* = 3.5, *p_corr_* = 0.018).

### 3.2 Cortical tracking of linguistic units

Figure 3 displays group-averaged SNR spectra and sensor distributions for each condition and age group. There was a clear peak of SNR at 2.5 Hz (i.e., word rate, *f*_word_) in all conditions and age groups, demonstrating excellent synchronization to the sequence of monosyllabic words. Other peaks were also noticeable at 1.25 Hz (i.e., phrase rate, *f*_phrase_) and at 0.625 Hz (i.e., sentence rate, *f*_sentence_), but only in the *Meaningful* and *Meaningful_noise_* conditions where sentential units were present. In all cases, SNR peaked in MEG sensors covering bilateral fronto-temporal areas.

**Figure 3.**
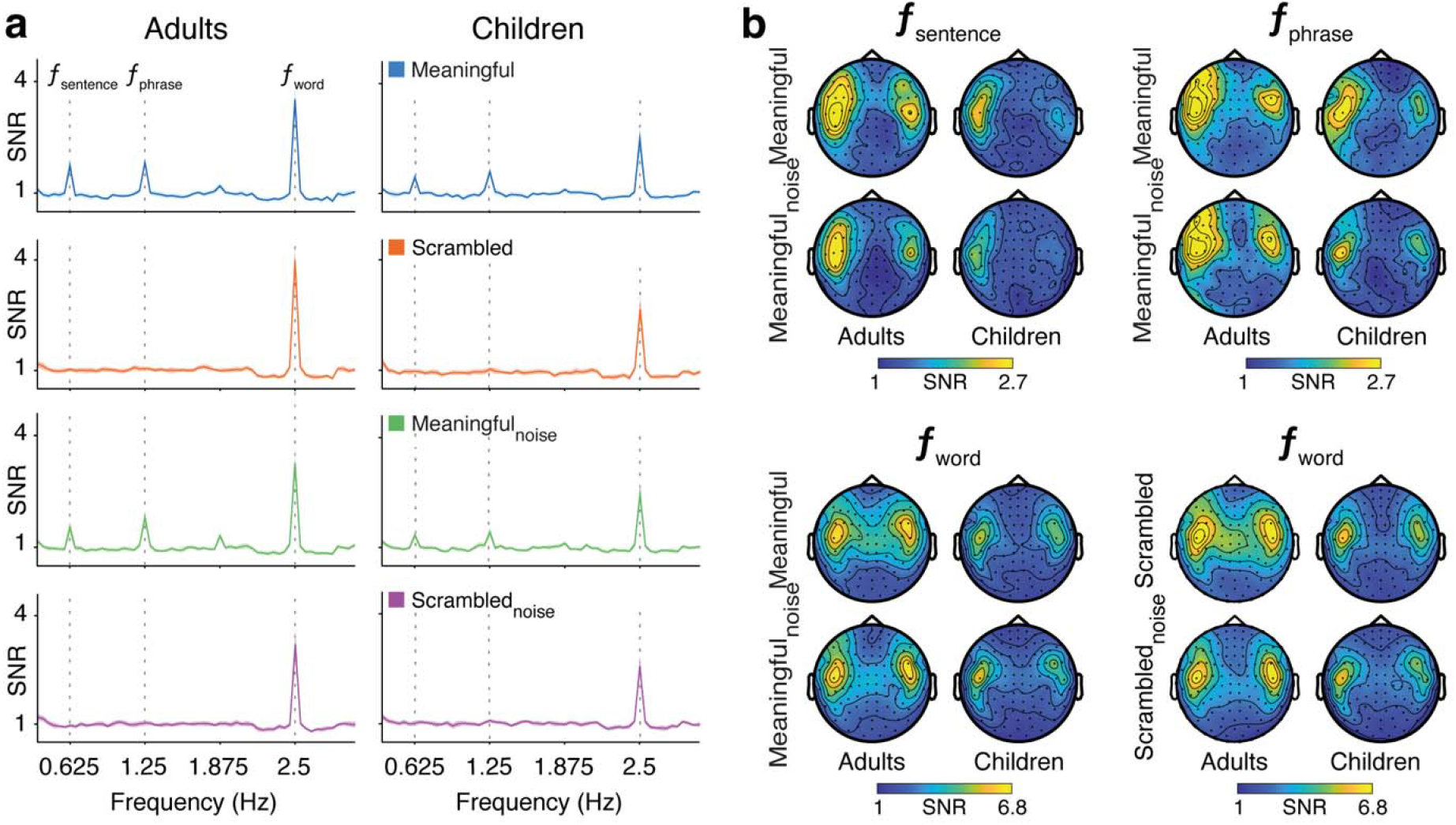
Group-level SNR spectra (a) and the corresponding topographical maps (b) for the three frequencies of interest and both groups.

Table 1 provides the number of adults and children showing significant SNR at *f*_word_, *f*_phrase_ and *f*_sentence_ for each condition. Tracking at *f*_word_ was significant in all participants and conditions. Furthermore, tracking at *f*_phrase_ and *f*_sentence_ were significant in most of the subjects in *Meaningful* and *Meaningful_noise_* conditions and in less than 3 subjects in the *Scrambled* and *Scrambled_noise_* conditions. Interestingly, sentence tracking (*f*_sentence_) in children was significant only in about half of the children.

**Table 1.**
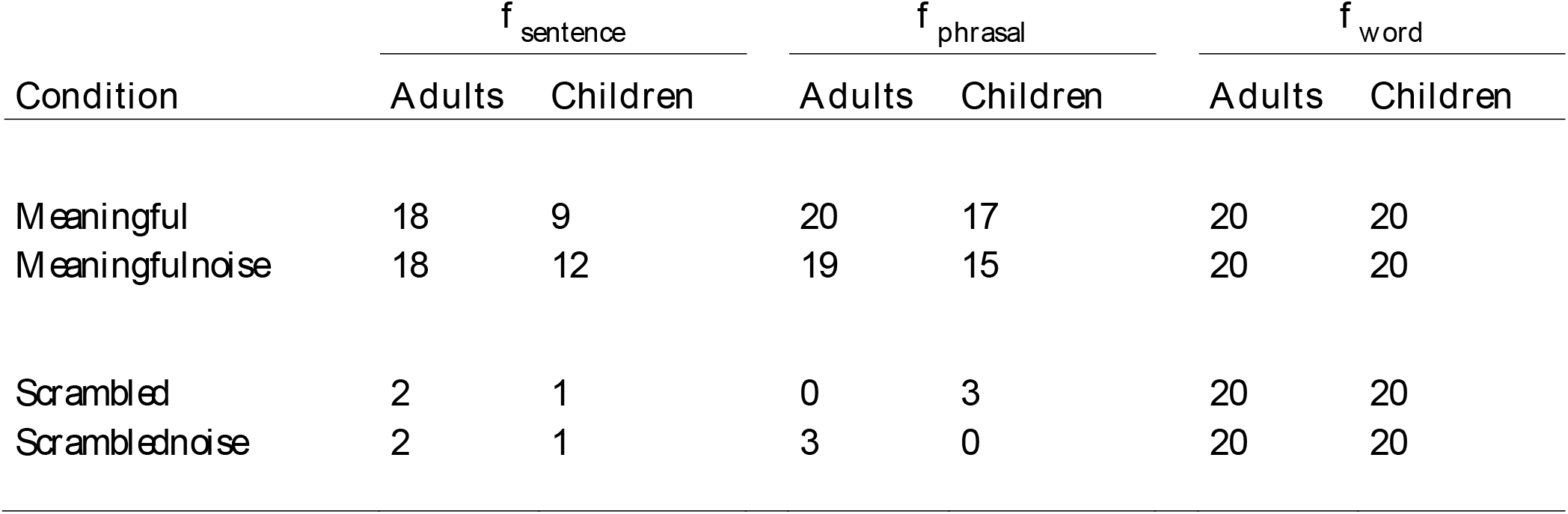
Number of adults and children showing statistically significant peaks in each frequency and condition.

| Condition | $f_{\text{sentence}}$ | | $f_{\text{phrasal}}$ | | $f_{\text{word}}$ | |
| --- | --- | --- | --- | --- | --- | --- |
|  | Adults | Children | Adults | Children | Adults | Children |
| *Meaningful* | 18 | 9 | 20 | 17 | 20 | 20 |
| *Meaningfulnoise* | 18 | 12 | 19 | 15 | 20 | 20 |
| *Scrambled* | 2 | 1 | 0 | 3 | 20 | 20 |
| *Scramblednoise* | 2 | 1 | 3 | 0 | 20 | 20 |

### 3.3 Cortical source localization

Figure 4 displays SNR source distribution in all conditions and in both age groups for each frequency of interest.

**Figure 4.**
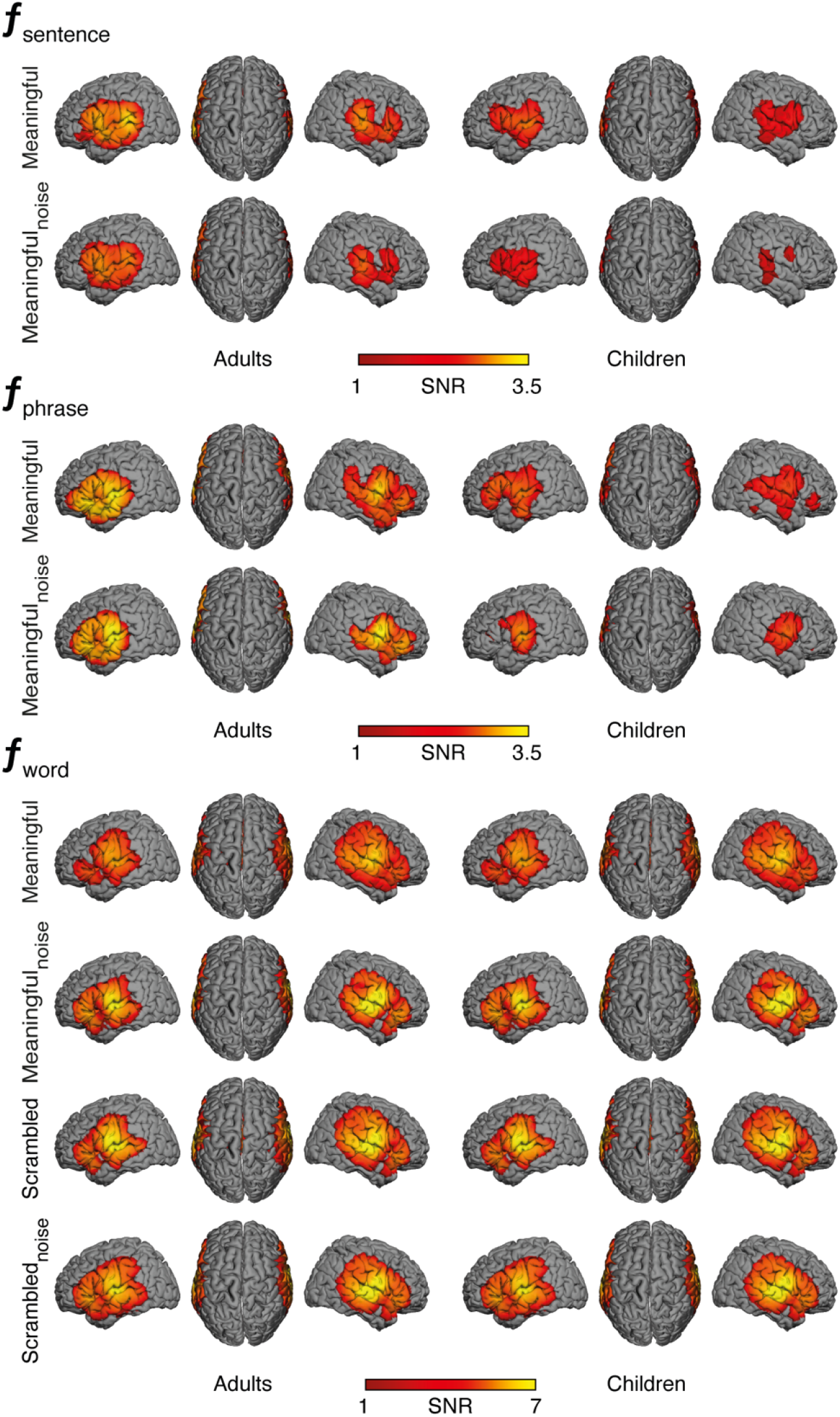
Source distributions of the SNR for each group, condition, and frequency of interest

Table 2 presents the coordinates of the significant local maxima of SNR. They peaked at auditory temporal areas and at inferior frontal gyrus bilaterally for all frequencies of interest. It is worth highlighting that it specifically peaked at bilateral pars opercularis at *f*_sentence_ but at bilateral pars orbitalis and triangularis at *f*_phrase_ and *f*_word_.

**Table 2.** Local maxima of group-level SNR map for all frequencies of interest: MNI coordinates and SNR. STAC = Supra-temporal auditory cortex; IFG = Inferior frontal gyrus; Op = Pars opercularis; Tri = Pars triangularis; Orb = Pars orbitalis.

| Frequency | Regions | x | y | z | SNR |
| --- | --- | --- | --- | --- | --- |
| $f_{\text{sentence}}$ | STAC | -50 | -25 | 8 | 2.69 |
|  |  | 56 | -22 | 12 | 2.25 |
|  | IFG Op | -42 | 9 | 14 | 2.62 |
|  |  | 43 | 12 | 8 | 1.93 |
| $f_{\text{phrasal}}$ | STAC | -43 | -13 | 6 | 3.51 |
|  |  | 57 | 0 | 12 | 2.92 |
|  | IFG Tri | -35 | 30 | 4 | 2.96 |
|  | IFG Orb | 51 | 37 | 6 | 2.35 |
| $f_{\text{word}}$ | STAC | -42 | -12 | 12 | 6.93 |
|  |  | 51 | -6 | 10 | 6.52 |
|  | IFG Tri | -40 | 29 | 1 | 4.38 |
|  |  | 45 | 32 | -4 | 4.19 |

### 3.4 Effect of age group, noise and hemisphere on the cortical tracking of hierarchical linguistic units

Figure 5 displays the SNR values in *Meaningful* and *Meaningful_noise_* conditions, in both age groups and both hemispheres, for each frequency for the two identified sources of CTS (i.e., inferior frontal gyri, **Figure 5a**, and auditory temporal area, **Figure 5b**).

**Figure 5.**
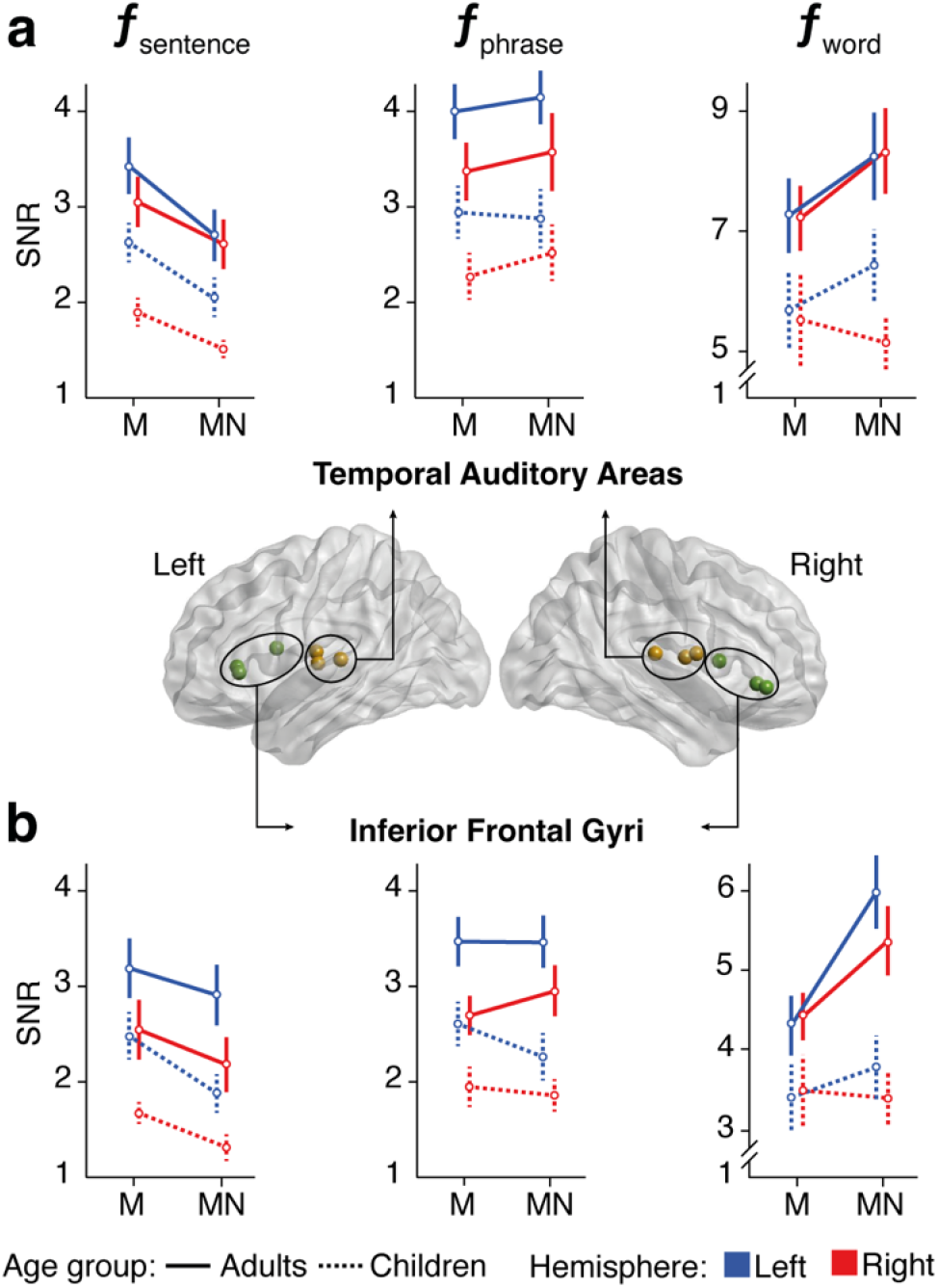
SNR of CTS sources (mean ± 2 SEM) in *Meaningful* (M) and *Meaningful_noise_* (MN) conditions, in both age groups and in hemispheres, for each frequency at the two regions of interest: (a) inferior frontal gyri and (b) auditory temporal areas. Significant local maxima are illustrated on the MNI glass brain.

Table 3 summarizes the results of the separate ANOVAs performed on SNR values in auditory temporal areas and inferior frontal gyri, at *f_sentence_*, *f_phrase_* and *f_word_* with factors noise, age group and hemisphere. The ANOVAs revealed a significant effect of age group for both cortical areas, explained by lower SNR values in children compared with adults. They also revealed a significant effect of the hemisphere at *f*_sentence_ and *f*_phrase_ but not at *f*_word_ in both cortical areas, reflecting higher SNR in the left hemisphere compared with the right. No significant interactions were found at *f*_sentence_ and *f*_phrase_. Analyses also revealed a main effect of noise at *f*_sentence_, reflecting higher SNR in *Meaningful* compared with *Meaningful_noise_* condition. A main effect of noise was also uncovered at *f*_word_ but only in the inferior frontal gyri, with significant interactions between noise and hemisphere, and between age and noise. These interactions were reflecting significantly higher SNR in *Meaningful_noise_* compared with *Meaningful* condition (1) in the left hemisphere (*t_39_* = 3.40, *p*_corr_ = 0.008) but not in the right hemisphere (*t_39_* = 1.42, *p*_corr_ = 0.98), and (2) in adults (*t_19_* = 3.39, *p*_corr_ = 0.0098) but not in children (*t_39_* = 0.36, *p*_corr_ = 1).

**Table 3.** Factors affecting the cortical tracking of hierarchical linguistic units. All significant results are in boldface.

| Region | Factor | $f_{\text{sentence}}$ | | $f_{\text{phrasal}}$ | | $f_{\text{word}}$ | |
| --- | --- | --- | --- | --- | --- | --- | --- |
| | | $F_{1,38}$ | $p$ | $F_{1,38}$ | $p$ | $F_{1,38}$ | $p$ |
| Temporal auditory areas | Age | **16.65** | **<0.0001** | **12.24** | **0.001** | **11.04** | **0.002** |
|  | Noise | **24.45** | **<0.0001** | 0.73 | 0.40 | 1.93 | 0.17 |
|  | Hemisphere | **5.99** | **0.019** | **7.00** | **0.012** | 0.71 | 0.41 |
|  | Age x Hemisphere | 1.34 | 0.25 | 0.039 | 0.85 | 0.80 | 0.38 |
|  | Age x Noise | 0.29 | 0.59 | 0.07 | 0.79 | 0.96 | 0.34 |
|  | Noise x Hemisphere | 2.56 | 0.12 | 0.67 | 0.42 | 1.15 | 0.29 |
|  | Age x Hemisphere x Noise | 0.02 | 0.88 | 0.33 | 0.57 | 1.87 | 0.18 |
| Inferior frontal gyri | Age | **11.27** | **0.002** | **17.09** | **<0.0001** | **12.38** | **<0.0001** |
|  | Noise | **12.40** | **0.001** | 0.13 | 0.72 | **7.05** | **0.012** |
|  | Hemisphere | **11.91** | **0.001** | **12.96** | **<0.0001** | 0.92 | 0.35 |
|  | Age x Hemisphere | 0.00 | 0.99 | 0.12 | 0.74 | 0.06 | 0.82 |
|  | Age x Noise | 0.46 | 0.50 | 1.78 | 0.19 | **4.60** | **0.04** |
|  | Noise x Hemisphere | 0.19 | 0.67 | 2.31 | 0.14 | **5.45** | **0.025** |
|  | Age x Hemisphere x Noise | 0.80 | 0.38 | <0.0001 | 0.98 | 0.24 | 0.63 |

## 4. Discussion

This study shows that the child brain tracks the hierarchical linguistic units of clear speech devoid of prosody, with a left-hemisphere dominance as the adult brain, but with reduced accuracy. This study also demonstrates that a multi-talker noise similarly reduces grammar-based cortical tracking of sentences in children and adults. Critically, when such noise is present, the adult brain increases the tracking of monosyllabic words while there is no evidence for such a mechanism in children.

### Reduced grammar-based cortical tracking of speech hierarchical linguistic units in children

In a quiet environment, adults and children exhibited a clear peak of SNR, indicative of the presence of CTS, at word frequency in all conditions, and at phrase and sentence frequencies only when words formed meaningful sentences. In both groups, grammar-based cortical tracking of words, phrases and sentences originated from inferior frontal gyri and temporal auditory areas, which are key nodes of the speech processing network (Hickok and Poeppel, 2007; Friederici, 2011) that have already been highlighted in studies investigating the cortical tracking of natural connected speech in adults and children (Vander Ghinst et al., 2019; Bertels et al., 2022).

CTS was nevertheless lower in children compared with adults for all linguistic units (i.e., words, phrases, sentences). Several non-exclusive hypotheses can be raised to explain this finding. First, these results might illustrate the key role prosodic cues (Mehler et al., 1988; Kalashnikova et al., 2018; Teixidó et al., 2018; Myers et al., 2019) play in childhood in supporting the cortical tracking of linguistic units in synergy with grammar-based knowledge (Mehler et al., 1988; Kalashnikova et al., 2018; Teixidó et al., 2018; Myers et al., 2019). Still, the reduced cortical tracking of monosyllabic words in *Meaningful* condition does not really support this hypothesis. Second, connected speech processing also involves attentional processes (Sanes and Woolley, 2011; Jones et al., 2015; Shinn-Cunningham et al., 2017; Thompson et al., 2017, 2019). A certain level of attention is indeed required for combining syllables into words (Ding et al., 2018), or words into phrases/sentences (Makov et al., 2017). Thus, the reduced CTS observed in children might also be driven by reduced attention that is known to develop through childhood (Leibold, 2012). Finally, this might be explained by reduced SNR of MEG signals in children due to smaller head size or increased head movements (Wehner et al., 2008; Witton et al., 2014), but this hypothesis is questionable. Indeed, similar levels of CTS were previously found during natural connected speech listening at the sentential rate (<1Hz) in children of similar age compared to adults (Vander Ghinst et al., 2019). The future use of on-scalp MEG based on optically pumped magnetoencephalography should clarify this issue (Hill et al., 2019; de Lange et al., 2021). Further studies are thus needed to address these issues and better understand the origin of the observed difference between children and adults.

### Impact of a multi-talker background on the cortical tracking of hierarchical linguistic units

Behavioral scores (SiN audiometry, word identification and intelligibility ratings) were significantly lower in children compared with adults in noise, while they were similar in a quiet environment. These results are perfectly in line with the reduced SiN processing abilities of children <10 years (Elliott, 1979; Hall et al., 2002).

In a multi-talker background, adults and children’s cortical activity tracked phrases and sentences, reflecting ongoing grammar-based neural building of the hierarchical linguistic units in such adverse auditory scene. This is in line with previous studies conducted in adults (Mesgarani and Chang, 2012; Zion Golumbic et al., 2013; O’Sullivan et al., 2015; Rimmele et al., 2015; Vander Ghinst et al., 2016; Fuglsang et al., 2017; Destoky et al., 2019) and children (Destoky et al., n.d.; Vander Ghinst et al., 2019; Bertels et al., 2022) showing significant cortical tracking of the attended speech in a multi-talker background at similar SNR (i.e., +3 to 0 dB). Critically, the amplitudes of auditory temporal and inferior frontal responses at sentence frequency were significantly lower in noisy conditions, which is comparable to the dampening of the cortical tracking of natural connected speech previously reported at ~0.5-Hz in children and adults when speech is polluted by a multi-talker background (Vander Ghinst et al., 2016, 2019; Giordano et al., 2017; Destoky et al., 2019, n.d.; Bertels et al., 2022). This noise-related reduction in the cortical tracking of the attended speech at the sentence level probably accounts for the lower behavioral scores observed both in children and adults. Still, no difference was observed in the effect of noise on the tracking of phrases/sentences between children and adults. But considering that children have a weaker tracking of phrases/sentences in the *Meaningful* condition compared with adults, the similar effect of noise between children and adults might actually have a higher functional impact in children that would partly explain why school-aged children have lower SiN processing abilities than adults.

The cortical tracking of phrases was not affected by noise. Binding two elements into a syntactic hierarchy is considered as the most basic operation of the hierarchic syntactic building (Mueller et al., 2012; Friederici, 2020) and has been shown to be already operational in prelinguistic infants (Mueller et al., 2012; Friederici, 2020). This might explain the robustness of the cortical tracking of phrases at this sound level of babble noise even in children.

Crucially, the noisy background induced an increase in the cortical tracking of monosyllabic words in the IFG, in adults but not in children. This result is reminiscent of what has been observed for the cortical tracking of natural connected speech at the syllabic rate (i.e, 4-8 Hz): while the cortical tracking of the attended speech was significantly increased by a multi-talker background (up to 0dB) in adults, such increase was not observed in children (<10 years) (Vander Ghinst et al., 2019) and in adults with impaired speech perception in noise and normal peripheral auditory function (ISPiN) (Vander Ghinst et al., 2021). As the strength of the CTS has been related to speech intelligibility (Peelle et al., 2013; Doelling et al., 2014), these findings highlight key neural correlates of the reduced behavioral abilities of school-aged children and adults with ISPiN to understand SiN (Vander Ghinst et al., 2019, 2021). Indeed, these data highly suggest that the ability to increase the neural tracking of (sub-)lexical linguistic units in adverse auditory scenes plays a key functional role in the human capacity to properly perceive and understand SiN. Impairments in this ability would represent a common neural correlate to physiological or pathological conditions characterized by a reduced SiN understanding at the behavioral level.

### Hemispheric dominance of the cortical tracking of speech hierarchical linguistic units

In the absence of any prosodic cue, the CTS in a quiet environment was clearly left-hemisphere dominant at the phrase and sentence frequencies, both in children and adults. Contrastingly, previous studies using natural connected speech revealed that the auditory system tracks the slow fluctuations of speech temporal envelope (< 2Hz) preferentially in the right hemisphere (Bourguignon et al., 2013, 2018; Gross et al., 2013; Molinaro et al., 2016; Vander Ghinst et al., 2016, 2019; Destoky et al., 2019). These opposing results therefore provide additional empirical evidence supporting the hypothesis that the right-dominant CTS is mainly driven by the neural tracking of prosodic cues (Friederici, 2002; Bourguignon et al., 2013).

The present study identified a similar impact of noise on the grammar-based cortical tracking of sentences in children and adults. Yet, previous studies performed in school-aged children and adults consistently demonstrated that the cortical tracking of attended natural connected speech at phrase/sentence levels is more robust to noise in the left hemisphere compared with the right (Peelle et al., 2013; Rimmele et al., 2015; Vander Ghinst et al., 2016, 2019; Destoky et al., 2019). The right-hemisphere CTS is also more easily corrupted by noise in children than in adults (Vander Ghinst et al., 2019; Bertels et al., 2022). Considering that right-hemisphere CTS appears to be mainly driven by the neural tracking of prosodic cues, these data might thus provide indirect evidence suggesting that children’s behavioral SiN processing difficulties might also be rooted in suboptimal non-verbal (i.e., prosodic) rather than grammar-based neural processing of the attended speech stream. Further studies are needed to confirm that hypothesis.

## 5. Conclusion

As compared to adults, school-aged children appear to be unable to enhance the cortical tracking of monosyllabic words in a multi-talker background noise. This might partly contribute to their lower behavioral ability to understand speech in noise. This effect comes in addition to a restricted cortical tracking of prosodic elements that has been previously shown in such adverse auditory scenes.

## Acknowledgments

Maxime Niesen and Marc Vander Ghinst were supported by the Fonds Erasme (Brussels, Belgium). Mathieu Bourguignon and Julie Bertels have been supported by the program Attract of Innoviris (grants 2015-BB2B-10 and 2019-BFB-110). Julie Bertels has been supported by a research grant from the Fonds de Soutien Marguerite-Marie Delacroix (Brussels, Belgium). Xavier De Tiège is Clinical Researcher at the Fonds de la Recherche Scientifique (FRS-FNRS, Brussels, Belgium).

This study and the MEG project at CUB Hôpital Erasme were financially supported by the Fonds Erasme (Research Convention: “Les Voies du Savoir”, Fonds Erasme, Brussels, Belgium).

## Data and code availability statement

MEG data used in this study will be made available upon reasonable request to the corresponding author.

